# Flexible expressed region analysis for RNA-seq with derfinder

**DOI:** 10.1101/015370

**Authors:** Leonardo Collado-Torres, Abhinav Nellore, Alyssa C. Frazee, Christopher Wilks, Michael I. Love, Ben Langmead, Rafael A. Irizarry, Jeffrey T. Leek, Andrew E. Jaffe

## Abstract

**Background:** Differential expression analysis of RNA sequencing (RNA-seq) data typically relies on reconstructing transcripts or counting reads that overlap known gene structures. We previously introduced an intermediate statistical approach called differentially expressed region (DER) finder that seeks to identify contiguous regions of the genome showing differential expression signal at single base resolution without relying on existing annotation or potentially inaccurate transcript assembly.

**Results:** We present the `derfinder` software that improves our annotation-agnostic approach to RNA-seq analysis by: (1) implementing a computationally efficient bump-hunting approach to identify DERs which permits genome-scale analyses in a large number of samples, (2) introducing a flexible statistical modeling framework, including multi-group and time-course analyses and (3) introducing a new set of data visualizations for expressed region analysis. We apply this approach to public RNA-seq data from the Genotype-Tissue Expression (GTEx) project and BrainSpan project to show that `derfinder` permits the analysis of hundreds of samples at base resolution in R, identifies expression outside of known gene boundaries and can be used to visualize expressed regions at base-resolution. In simulations our base resolution approaches enable discovery in the presence of incomplete annotation and is nearly as powerful as feature-level methods when the annotation is complete.

**Conclusions:** `derfinder` analysis using expressed region-level and single base-level approaches provides a compromise between full transcript reconstruction and feature-level analysis.

The package is available from `Bioconductor` at www.bioconductor.org/packages/derfinder.

## 1 Introduction

The increased flexibility of RNA sequencing (RNA-seq) has made it possible to characterize the transcriptomes of a diverse range of experimental systems, including human tissues [1, 2, 3], cell lines [4, 5] and model organisms [6, 7]. The goal of many experiments involves identifying differential expression with respect to disease, development, or treatment. In experiments using RNA-seq, RNA is sequenced to generate short “reads” (36-200+ base pairs). These reads are aligned to a reference genome, and this alignment information is used to quantify the transcriptional activity of both annotated (present in databases like Ensembl) and novel transcripts and genes.

The ability to quantitatively measure expression levels in regions not previously annotated in gene databases, particularly in tissues or cell types that are difficult to ascertain, is one key advantage of RNA-seq over hybridization-based assays like microarray technologies. As complicated transcript structures are difficult to completely characterize using short read sequencing technologies [8], the most mature statistical methods used for RNA-seq analysis rely on existing annotation for defining regions of interest - such as genes or exons - and counting reads that overlap those regions [9]. These counts are then used as measures of gene expression abundance for downstream differential expression analysis [10, 11, 12, 13, 14, 15, 16, 17, 18]. Unfortunately, the gene annotation may be incorrect or incomplete, which can affect downstream modeling of the number of reads that cross these defined features.

We previously proposed an alternative statistical model for finding differentially expressed regions (DERs) that first identifies regions that show differential expression signal and then annotates these regions using previously annotated genomic features [19]. This analysis framework first proposed using coverage tracks (i.e. the number of reads aligned to each base in the genome) to identify differential expression signal at each individual base and merges adjacent bases with similar signal into candidate regions. However, the software for our first version was limited to small sample sizes, the ability to interrogate targeted genomic loci, and comparisons between only two groups.

Here we expand the DER finder framework to permit the analysis of larger sample sizes with more flexible statistical models across the genome. This paper introduces a comprehensive software package called derfinder built upon base-resolution analysis, which performs coverage calculation, preprocessing, statistical modeling, region annotation and data visualization. This software permits differential expression analysis at both the single base level, resulting in direct calculation of DERs [20], and a feature summarization we introduce here call “expressed region” (ER)-level analysis. We show that ER analysis allows us to perform base resolution analysis on larger scale RNA-seq data sets using the BrainSpan project [21] http://developinghumanbrain.org and Genotype-Tissue Expression (GTEx) project data [3] to demonstrate that `derfinder` can identify differential expression signal in regions outside of known annotation without assembly. We use these DERs to illustrate the post-discovery annotation capabilities of `derfinder` and label each DER as exonic, intronic, intergenic or some combination of those labels. We show that some of these DERs we identify are outside of annotated protein coding regions and would not have been identified using gene or exon counting approaches.

In the GTEx data, we identify differentially expressed regions (DERs) that differentiate heart (left ventricle), testis and liver tissues for 8 subjects. There are many potential reasons for this observed intronic expression including intron retention, background levels of mis-transcription, or incomplete protein-coding annotation. A subset of these strictly intronic ERs are associated with tissue differences, even conditional on the expression of the nearest annotated protein-coding region. However, we point out that intronic expression may be artifactual and it our package permits visualization and discovery of potential expression artifacts not possible with other packages.

Finally, using simulated differentially expressed transcripts, we demonstrate that when transcript annotation is correct, `derfinder` is nearly as powerful as exon-count based approaches with statistical tests performed by `DESeq2` [14] (or `edgeR`-robust [13]) and `ballgown` [22] after summarizing the information using Rsubread [13] and `StringTie` [23] respectively. Finally, we also demonstrate that when annotation is incomplete, `derfinder` can be substantially more powerful than methods that rely on a complete annotation.

## 2 Materials & Methods

### 2.1 Overview of R Implementation

We chose to implement `derfinder` entirely in the R statistical environment www.R-project.org/. Our software includes upstream pre-processing of BAM and/or BigWig files into base-resolution coverage. At this stage the user can choose to summarize the base resolution coverage into feature-level counts and apply popular feature-level RNA-seq differential expression analysis tools like `DESeq2` [14], `edgeR`-robust [13], *limma* [16, 15] and *voom* [17].

`derfinder` can be used to identify regions of differential expression agnostic to existing annotation. This can be done with either the expressed regions (ER)-level or single base-level approaches, described in detail in the following subsection and Supplementary Section 2.1. The resulting regions can then be visualized to identify novel regions and filter out potential artifacts.

After differential expression analysis, `derfinder` can plot DERs using base-resolution coverage data by accessing the raw reads within differentially expressed regions for posthoc analysis like clustering and sensitivity analyses. We have also created a lightweight annotation function for quickly annotating DERs based on existing transcriptome annotation, including the UCSC knownGene hg19, Ensembl p12, and Gencode v19 databases as well as newer versions.

Vignettes with detailed instructions and examples are available through the Bioconductor pages for `derfinder` and `derfinderPlot`. The main functions for the expressed region and single base-level approaches are further described in Supplementary Section 1.1.

### 2.2 Expressed region level analysis

In the expressed region approach, we compute the mean coverage for all base pairs from all the samples and filter out those below a user specified cutoff. Contiguous bases passing this filtering step are then considered a candidate region (Figure 2A). Then for each sample, we sum the base-level coverage for each such region in order to create an expression matrix with one row per region and one column per sample. This matrix can then be used with feature-level RNA-seq differential expression analysis tools.

**Figure 1.**
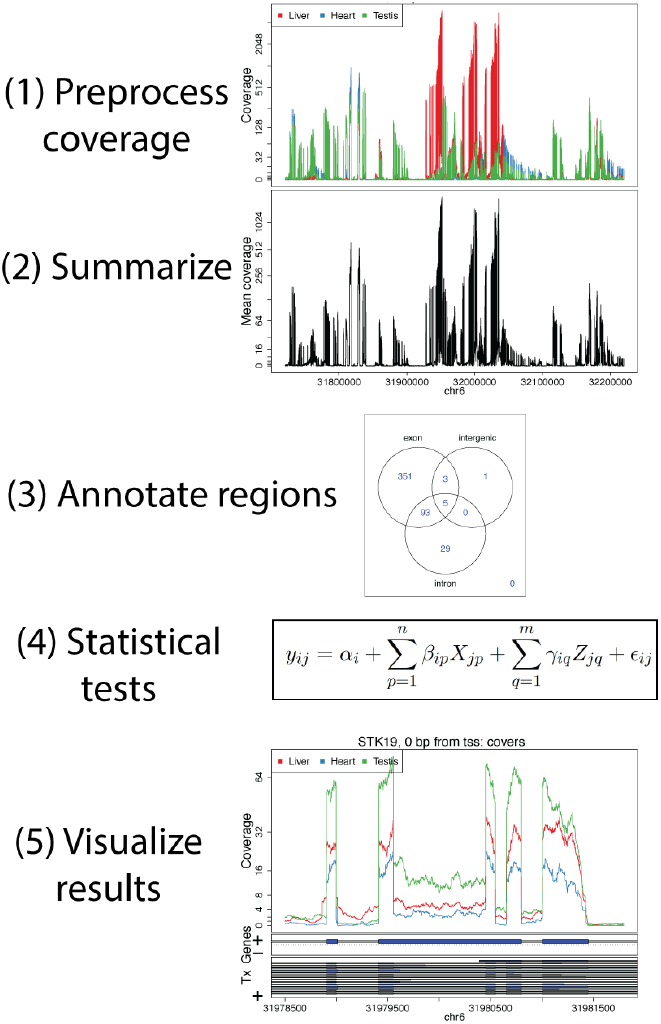
An overview of the derfinder suite. The `derfinder` software package includes functions for processing and normalizing coverage per sample, performing statistical tests to identify differentially expressed regions, labeling those regions with known annotation, and visualizing the results across groups.

**Figure 2.**
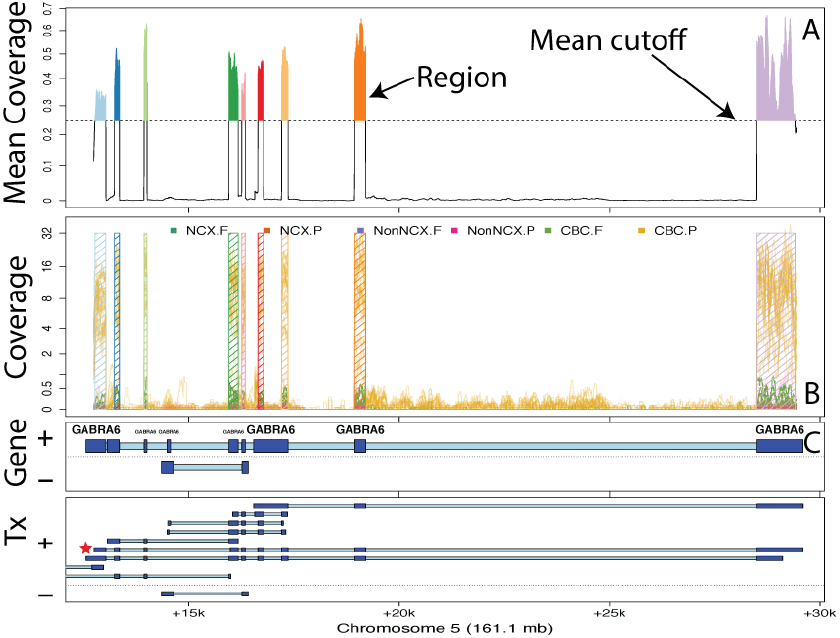
Finding regions via expressed region-level approach on chromosome 5 with *BrainSpan* data set. **A** Mean coverage with segments passing the mean cutoff (0.25) marked as regions. **B** Raw coverage curves superimposed with the candidate regions. Coverage curves are colored by brain region and developmental stage (NCX: Neocortex: Non-NCX: Non-neocortex, CBC: cerebellum, F: fetal, P: postnatal). **C** Known exons (dark blue) and introns (light blue) by strand for genes and subsequent transcripts in the locus. The DERs best support the *GABRA6* transcript with a red star, indicating the presence of a differentially expressed transcript.

### 2.3 Annotation and “Genomic State” Objects

We have implemented a “genomic state” framework to efficiently annotate and summarize resulting regions, which assigns each base in the genome to exactly one state: exonic, intronic, or intergenic, based on any existing or user-defined annotation (e.g. UCSC, Ensembl, Gencode). At each base, we prioritize exon > intron > unannotated across all annotated transcripts.

Overlapping exons of different lengths belonging to different transcripts are reduced into a single “exonic” region, while retaining merged transcript annotations. We have a second implementation that further defines promoters and divides exonic regions into coding and untranslated regions (UTRs) which may be useful for the user to more specifically annotate regions - this implementation prioritizes coding exon > UTR > promoter > intron > unannotated.

### 2.4 Data Processing for Results in Main Manuscript

#### 2.4.1 BrainSpan data

BigWig files for all 487 samples across 16 brain regions were downloaded from the *BrainSpan* website [21]. The samples for *HSB169.A1C, HSB168.V1C* and *HSB168.DFC* were dropped due to quality issues. Based on exploratory analyses the coverage was assumed to be reads-per-million mapped reads in this data set. We set the coverage filter to 0.25 for both the single base-level and ER-level `derfinder` approaches. Since the coverage is already adjusted to reads per million mapped reads we did not include a library size adjustment term in the single base-level `derfinder` analysis (see Supplementary Section 2.1 for details on this adjustment term). The details for the single base-level `derfinder` analysis are described further in Supplementary Section 2.2. For the ER-level approach we only considered regions longer than 5 base-pairs.

We sought to identify differences in expression across brain region (neocortical regions: DFC, VFC, MFC, OFC, M1C, S1C, IPC, A1C, STC, ITC, V1C and non-neocortical regions: HIP, AMY, STR, MD, and CBC) and developmental stage (fetal versus postnatal). We therefore fit the following region-by-stage interaction alternative model, which included main effects for fetal versus postnatal (binary) and categorical brain region variable (15 region indicators, relative to A1C), and interaction terms for each brain region and developmental stage. This resulted in a total of 32 terms in the model (intercept; 16 main effects, 15 interaction terms).

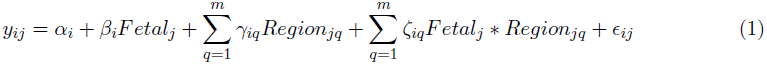

We compared the above model to an intercept-only model where *y_ij_* is the log (mean base-level coverage +1) using the `lmFit` function from `limma` [16, 15]. The p-values for the ER-level DERs were adjusted via the Bonferroni method and those with adjusted p-values less than 0.05 were determined to be significant. We then calculated the mean coverage for each significant expressed region DERs in each sample, resulting in a mean coverage matrix (DERs by samples), and we performed principal component analysis (PCA) on this log2-transformed matrix (after adding an offset of 1), which were subsequently plotted in Figure 5.

**Figure 3.**
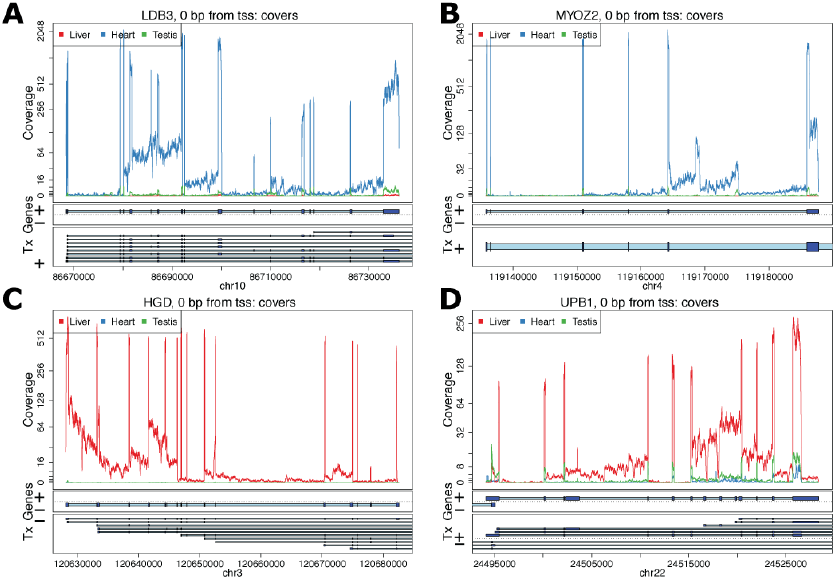
Figure 3: Coverage plots for the average coverage levels for the GTEx example. Average coverage profile for heart (blue), liver (red), and testis (green) from the GTEx example near genes: **A** *LDB3*, **B** *MYOZ2*, **C** *HGD*, and **D** *UPB1*.

#### 2.4.2 GTEx data

We selected samples from individuals that had data from heart (left ventricle), liver and testis tissues with RIN values greater than 7. 8 subjects matched this criteria and we selected only 1 sample if their tissue was analyzed more than once, leaving us with 24 samples. The data was aligned using `Rail-RNA` [24] version 0.2.1 with the code as described at github.com/nellore/runs. We created a normalized mean BigWig file for these 4 samples adjusted for library sizes of 40 million reads. We then identified the ERs using a cutoff of 5 using the function `railMatrix` from `derfinder` version 1.5.19.

For each expressed region greater than 9bp, we assigned its annotation status by using a genomic state object created with the Ensembl GRCh38.p5 database. We then performed principal component analysis (PCA) on the log_2_-transformed matrix (after adding an offset of 1) separately for strictly exonic and strictly intronic ERs. Using `limma` [16, 15] functions `lmFit`, `ebayes` we fit an intercept-only null model and an alternative model with coefficients for tissue differences. For each ER we calculated a F-statistic and determined whether it was differentially expressed by tissue using a Bonferroni adjusted p-value cutoff of 0.05.

For the conditional expression analysis, we found the nearest exonic ER for each intronic ER using the `distanceToNearest` function from `GenomicRanges` [25]. For each intronic ER we fitted two linear regression models for the log_2_-transformed coverage matrix (after adding an offset of 1). For the alternative model we used as covariates two tissue indicator variables (Heart as the reference) and the coverage from the nearest stricly exonic ER as shown in Equation (2) for ER *i* and sample *j*. For the null model we only used the coverage from the nearest exonic ER. We calculated an F-statistic using the `anova` function that tests whether *β*_1*i*_ or *β*_2*i*_ are equal to 0 and used a Bonferroni adjusted p-value cutoff of 0.05 to identify which intronic ERs had differential expression adjusting for the coverage at the nearest exonic ER.

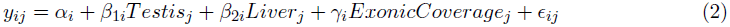

#### 2.4.3 Simulated data

We simulated 100 bp paired-end reads (250bp fragments, sd = 25) with polyester [26] for two groups with five samples per group from human chromosome 17 with uniform error rate of 0.005 and replicated this process three times. One sixth of the transcripts were set to have higher expression (2x) in group 2, a sixth to have lower expression in group 2 (1/2x) and the remaining two thirds to be equally expressed in both groups. Given a RNA-seq experiment with 40 million paired-end reads, assuming that all transcripts are equally expressed we would expect 1,989,247 of them to be from chromosome 17 based on the length of all exons using the known transcripts UCSC knownGene hg19 annotation. We used this information and the transcript length to assign the number of reads per transcript in chromosome 17 and generated the number of reads with the NB function from polyester with mean *μ* and size (see `stats::rnbinom` function in R) equal to ⅓*μ*. This resulted in an average of 2,073,682 paired-end reads per sample. For each simulation replicate, paired-end reads were aligned to to the hg19 reference genome using `HISAT` version 0.1.6-beta [27] and `Rail-RNA` version 0.2.2b [24]. We created a GTF file using all known transcripts from chromosome 17 as well as one with 20% of the transcripts missing (8.28% of exons missing). Using these two GTF files we performed transcript quantification with `StringTie` version 1.2.1 [23] as well as exon counting allowing multiple overlaps with the `featureCounts` function from `Rsubread` version 1.21.4 [13]. ERs were determined with `derfinder` version 1.5.19 functions `regionMatrix` and `railMatrix` respectively from the `HISAT` BAM and `Rail-RNA` BigWig output using a mean cutoff of 5 for libraries adjusted to 80 million single-end reads. Count matrices resulting for `featureCounts` and `derfinder` were analyzed with *DESeq2* [14] and *edgeR*-robust [18] controlling the FDR at 5% and testing for differences between the two groups of samples. We used `ballgown` version 2.2.0 [22] to perform differential expression tests using coverage at the transcript and exon levels, controlling the FDR at 5%.

The 3900 transcripts from chromosome 17 are composed in total by 39,338 exons (15,033 unique). To avoid ambiguous truth assignments, we used only the 3,868 that overlap only 1 transcript and assigned the truth status based on whether that transcript was set to have a high or low expression on group 2 for the replication replicate under evaluation. We assessed the different pipelines by checking if these 3,868 exons overlapped at least one differentially expressed unit: exons (`featureCounts` and `ballgown`), transcripts (`ballgown`), and ERs (`derfinder`) respectively. We then calculated the empirical power, false discovery rate and false positive rate.

## 3 Results

### 3.1 Overview of the derfinder package

The `derfinder` package includes functions for several stages in the analysis of data from an RNA-sequencing experiment (Figure 1).

First, `derfinder` includes functions for pre-processing coverage data from BAM files or bigWig coverage files. The base-level coverage data for multiple samples can be loaded and filtered since most bases will show zero or very low coverage across most samples. Then, the software allows for definition of contiguous regions that show average coverage levels above a certain threshold. These expressed regions are non-overlapping subsets of the genome that can then be counted to arrive at a matrix with an expression value for each region in each sample. Alternatively, the software provides options for counting exons or genes for use in more standard analysis pipelines.

Next, `derfinder` can be used to perform statistical tests on the region level expression matrix. These tests can be carried out using any standard package for differential expression of RNA-seq data including `edgeR` [10, 12], `DESeq` [11], `DESeq2` [14], or `limma`-voom [17]. In the analysis that follows we focus on using the `limma` package [16, 15].

`derfinder` can then be used to annotate the differentially expressed regions (DERs). We have developed functions that label each region according to whether it falls entirely in a previously annotated protein coding exon (exonic), entirely inside a previously annotated intronic region (intronic), or outside of any previously annotated gene (intragenic). The software also will report any region that overlaps any combination of those types of regions.

Finally, data from an expressed region analysis can be visualized using different visualization approaches. While region-level summaries can be plotted versus known phenotypes, `derfinder` also provides functions to plot base resolution coverage tracks for multiple samples, labeled with color according to phenotype.

We now provide more detail on each of these steps.

### 3.2 Finding expressed regions

The first step in a `derfinder` analysis is to identify expressed regions. Reads should be aligned using any splicing aware alignment tool such as `TopHat2` [28], `HISAT` [27] or `Rail-RNA` [24].

Base resolution coverage information can be read directly from the BAM files that are produced by most alignment software [28, 27, 24]. This process can be parallelized across multiple cores to reduce computational time. An alternative is to read bigWig [29] coverage files. Recent alignment software such as `Rail-RNA` [24] produces these files directly, or they can be created using sam-tools [30] or produced using the `derfinder` package. Reading bigWig files can produce significant computational and memory advantages over reading from BAM files.

The coverage information represents the number of reads that covers each genomic base in each sample. `derfinder` first filters out bases that show low levels of expression across all samples. Since most genomic bases are not expressed, this filtering step can reduce the number of bases that must be analyzed by up to 90%, reducing both CPU and memory usage. We originally proposed performing a statistical test for every base in the genome [19] and this approach is still supported by the `derfinder` package for backwards compatibility (Supplementary Section 1.2).

Here we focus on a new approach based on the bump-hunting methodology for region level genomic analysis [31] (Figure 2). This approach first calculates expressed regions (ERs) across the set of observed samples. For each base, the average, potentially library size-adjusted, coverage is calculated across all samples in the data set. This generates a vector of (normalized) mean level expression measurements across the genome. Then an average-coverage cutoff is applied to this mean coverage vector to identify bases that show minimum levels of expression. An expressed region is any contiguous set of bases that has expression above the mean expression cutoff.

The next step is to count the number of reads (including fractions of reads) that overlap each expressed region. As we have pointed out previously [19] that counting expression in genes and exons is complicated by overlapping annotation. Expressed regions are non-overlapping, so this means that each read can be unambiguously assigned to the appropriate region.

### 3.3 Expressed region level statistical tests

The result of the expressed region (ER) step is a coverage matrix with each row corresponding to one ER and each column corresponding to one sample. This count matrix can then be analyzed using statistical models that have been developed for gene or exon counts such as `limma` [16, 15], voom [17], `edgeR`-robust [18], and `DESeq2` [14]. We emphasize that unlike other feature-level counting approaches, our approach is annotation-agnostic: ERs are defined empirically using the observed sample data and coverage threshold. So if there is sufficient expression in a region outside of previously annotated genes it will be quantified and analyzed with our approach.

### 3.4 Visualizing differentially expressed regions

After statistical modeling, `derfinder` produces a set of DERs with summary statistics per region. They are stored as a `GRanges` object [25] and can be visualized with a range of packages from the Bioconductor suite. We have also developed several visualization tools specific to the `derfinder` approach.

These plots can be made at different levels of summarization. First, the `derfinder` and `derfinderPlot` packages provide a range of visualizations of coverage tracks at single base resolution. These plots can be used to identify coverage patterns that may diverge from annotated protein-coding regions. For example, using the GTEx example we can visualize genes that have consistently high intronic expression as shown in Figure 3. We show several examples of genes known to be functionally important in heart - *LBD3* and *MYOZ2* (Figure 3A,B) [32, 33], and liver - *HGD* and *UPB1* (Figure 3C,D) [34, 35]. The coverage profiles can provide additional insight into transcription, and well as potential technical artifacts, beyond the level of annotated genes, exons and transcripts, which we include in our base-resolution plots.

DERs can be grouped into larger regions by distance, which can be useful to identify potentially systematic artifacts such as coverage dips (Figure 4), perhaps due to sequence composition. Visualizing the base-level coverage for a set of nearby candidate DERs can reveal patterns that explain why one DER is sometimes fragmented into two or more shorter DERs. Coverage dips (Figure 4), spikes and data quality in general can affect the borders of the candidate DERs. Some artifacts can be discarded, like candidate DERs inside repetitive regions. Base-pairs inside repetitive regions available in repeat masker tracks can be flagged and filtered out from the analysis. Other known potentially problematic regions of the genome, like those with extreme GC content or mappability issues can also be filtered out, either before identifying candidate DERs or post-hoc.

**Figure 4.**
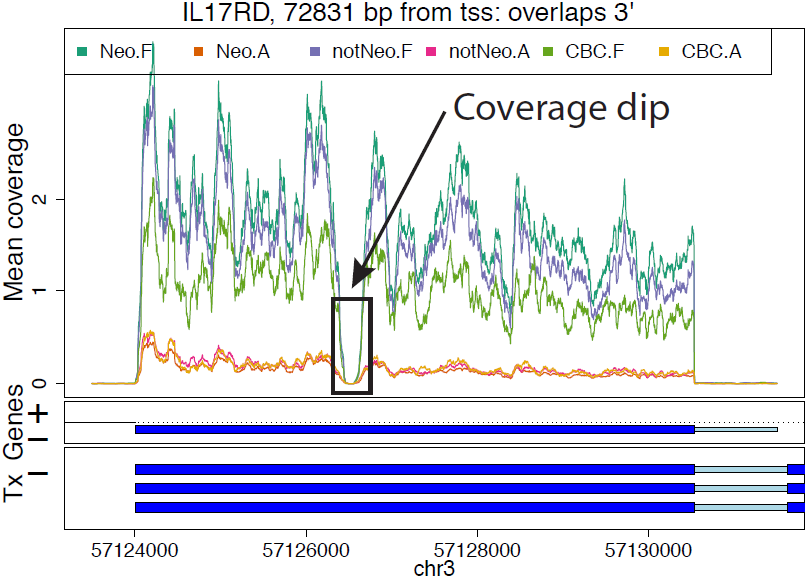
Example of a coverage dip. Mean coverage per group for the *BrainSpan* data set for a region that results in two DERs for a single exon due to a coverage dip. The genome segment shown corresponds to the DERs cluster ranked 15th in terms of overall signal by the single base-level approach applied to the *BrainSpan* data set.

**Figure 5.**
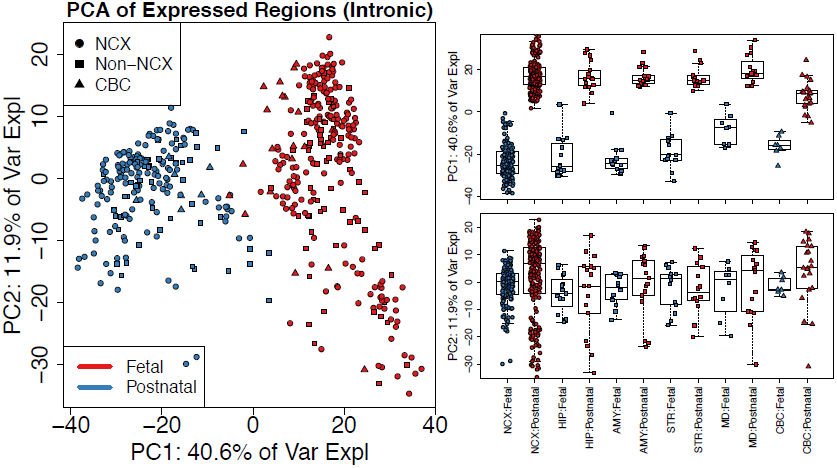
Principal components analysis reveals clusters of samples in the BrainSpan data set. (Left) First two principal components (PCs) with samples colored by sample type (F: Fetal or P: Postnatal) and shape given by brain region using only the strictly intronic expressed regions (ERs). Analysis of other subsets of ERs produce similar results (Supplementary Figure S2). (Right) Boxplots for PCs 1 and 2 by brain region (NCX: neocortex, HIP: hippocampus, AMY: amygdala, STR: striatum, MD: thalamus, CBC: cerebellum) and sample type with non-neocortex brain decomposed into its specific regions. Supplementary Figure S3 shows similar results using the single base-level approach.

### 3.5 Annotating differentially expressed regions

The DERs can be annotated to their nearest gene or known feature using `bumphunter` [31]. The basic approach is to overlap DERs genomic coordinates with the genomic coordinates of known genomic features. By default, `derfinder` labels each identified region as exonic, intronic, intragenic or some combination of those three labels.

A region may overlap multiple genomic features (say an exon and the adjacent intron). Using this information candidate DERs can further be compared to known gene annotation tables (Methods Section 2.3) to identify potentially novel transcription events. Using this information, visualizations of specific loci for overlap with annotation can be made with `derfinderPlot`. The regions can be exported to CSV files or other file formats for followup and downstream analyses. We have also developed a complementary R package for creating reproducible reports incorporating the annotation and visualization steps of the `derfinder` pipeline called `regionReport` [36].

### 3.6 Application: large-scale expression analysis at base resolution

We used `derfinder` to detect regions that were differentially expressed across the lifespan in the human brain. We applied `derfinder` to the *BrainSpan* RNA-seq coverage data (Methods Section 2.4.1), a publicly available data set consisting of 484 postmortem samples across 16 brain regions from 40 unique individuals that collectively span the full course of human brain development [21]. We used the expressed region approach described above for this analysis. For comparison we applied the single-based resolution approach previously utilized on independent dorsolateral prefrontal cortex RNA-seq data [20] (Supplementary Section 1.4).

We identified 174,610 ERs across the 484 samples with mean across-sample normalized coverage > 0.25, which constituted 34.57 megabases of expressed sequence. The majority (81.7%) of these ERs were labeled as strictly exonic while only a small subset (5.4%) were strictly non-exonic by Ensembl annotation. These ERs largely distinguished the fetal and postnatal samples using PCA - the first principal component explained 40.6% of the variance of the mean coverage levels and separated these developmental stages across all brain regions. This separation was consistent regardless of the annotation status of the DERs including in the strictly intronic regions (Figure 5 and Supplementary Figure S2). The separation between brain regions in intronic regions may be due to noisy or incorrect splicing [37] or may be due to missing annotation [19] or mistaken sequencing of pre-mRNA. The base resolution visualizations available as part of `derfinder` and `derfinderPlot` make it possible to explore to determine if it is biology or artifacts driving these expression differences.

The PCA plots also appear to show patterns consistent with potential artifacts such as batch effects [38] (Figure 5). Regardless, the new ER approach we present here provides options for analysts who wish to discover patterns of expression outside of known annotation on hundreds of samples - an analysis of this scope and scale was unfeasible with earlier versions of our single base resolution software [19].

Using statistical models where expression levels were associated with developmental stage (fetal versus postnatal) and/or brain region (Methods Section 2.4.1), we found that 129,278 ERs (74%) were differentially expressed by brain region and/or developmental stage at the ER-level controlling the family-wise error rate (FWER) at < 5% via Bonferroni correction. The 129,278 ER-level DERs overlapped a total of 17,525 Ensembl genes (13,016 with gene symbols), representing a large portion of the known transcriptome. Of the significant ER-level DERs, 93,355 (72.2%) overlapped at least 1 significant single base-level DER (Supplementary Section 1.4). Lack of overlap results from almost half (45.2%) of single base-level DERs having an average coverage lower than the expression cutoff determining ERs (0.25). For example, there was high expression only in the samples from a few brain regions, or only one development period. Decreasing the cutoff that defines the ERs from 0.25 to 0.1 results in a larger number of regions (217,085) that have a higher proportion of non-exonic sequence (12.1%), suggesting that the choice of this expression cutoff requires some initial exploratory data analysis.

Lastly, we highlight the utility of the ER-level analysis (using the original 0.25 cutoff) to identify regions differentially expressed within subsets of the data by analyzing brain regions within a single developmental period. We identified 1,170 ERs that were differentially expressed comparing striatum versus hippocampus samples in the fetal developmental stage. These DERs mapped to 293 unique genes. Genes more highly expressed in the striatum include *ARPP-21*, previously shown to localize in the basal ganglia [39], and dopamine receptor genes *DRD1* and *DRD2* [40]. Genes more highly expressed in the hippocampus in fetal life were strongly enriched for neurodevelopmental genes including *FZD7* [41], *ZBTB18* [42], and *NEUROD1* [43]. The ER-level analysis therefore permits subgroup analysis without the need to rerun the full `derfinder` single base-level pipeline - another improvement over previous versions of single base resolution analysis software [19].

### 3.7 Identification of expressed regions that differentiate tissues using a subset of the GTEx data

We selected a subset of subjects from the GTEx project [3] that had RNA-seq data from heart (left ventricle), liver and testis, specifically the eight subjects with samples that had RNA Integrity Numbers (RINs) greater than 7, given RIN’s impact on transcript quantification [44]. Using only one sequencing library from each subject aligned with `Rail-RNA` [24], we applied the ER-level `derfinder` approach with a cutoff of 5 normalized reads (after normalizing coverage to libraries of 40 million reads). We found a total of 163,674 ERs with lengths greater than 9 base-pairs. Figure 6A shows that 118,795 (72.6%) of the ERs only overlapped known exonic regions of the genome using the Ensembl GRCh38.p5 database [45].

we performed PCA on the log_2_ adjusted coverage matrix using just the 118,795 strictly exonic ERs (Figure 6B). Here the first two PCs explain 56.8% and 21.6% of the variance respectively and show three distinct clusters of samples that correspond to the tissue of the sample. We found that the 16,985 (10.4%) ERs (Figure 6A) that only overlap annotated introns can also differentiate tissues using PCA, as shown in Figure 6C. The total percent of variance explained by the first two principal components is slightly lower (44.4% + 26.6% = 71% versus 56.8% + 21.6% = 78.4%) when using only the strictly intronic ERs versus the strictly exonic ERs. This may represent a different biological signal and/or potentially noisy splicing (as in Figure 3B). but we use this example to illustrate the potential to use `derfinder` to explore regions outside of known annotation.

**Figure 6.**
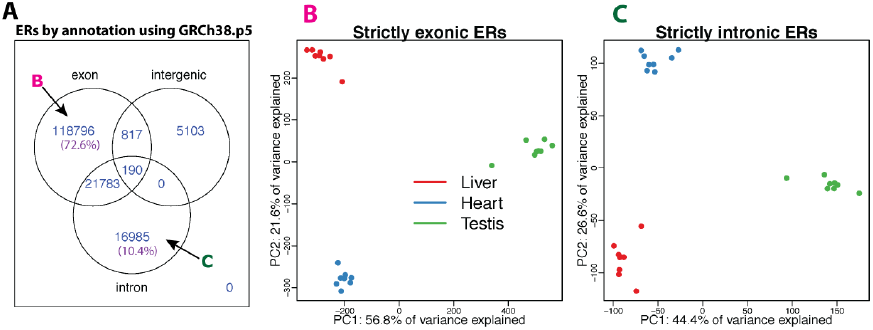
GTEx expressed regions analysis using 24 samples from the heart (left ventricle), liver and testis for 8 subjects. **A** expressed regions (longer than 9 bp) overlapping known annotation based on GRGh38.p5 (hg38). 72.6% of the ERs only overlap known exons (strictly exonic) while 10.4% only overlap known introns (strictly intronic). **B** First two principal components (PCs) with samples colored by sample type (red: liver, blue: heart, green: testis) using only the strictly exonic ERs. **C** First two PCs with samples colored by sample type using only only the strictly intronic ERs. The sign change of the second principal component is simply a rotation and the results are consistent between the strictly exonic and strictly intronic ERs.

Using `limma` [16, 15] to test for differential expression between tissues (Methods Section 2.4.2) we found that 42,880 (36.1%) of the strictly exonic ERs and 4,401 (25.9%) of the strictly intronic ERs were differentially expressed (FWER of 5% via Bonferroni correction). Overall 59,776 (36.5%) of the ERs were differentially expressed between tissues. Given the similar global patterns of expression between annotated and unannotated ERs, we considered the scenario that the strictly intronic ERs were differentially expressed between tissues in the same pattern as the nearest exonic ERs due to possible run-off transcription events. To assess this scenario we fitted a conditional regression for each strictly intronic ER adjusting for the coverage of the nearest strictly exonic ER. 749 (4.4%) of the strictly intronic ERs differentiate tissues while adjusting for the coverage at the nearest exonic ER at a FWER of 5%. Figure 7A,B shows an example where the expression is similar between tissues in the nearest exonic ER but there is a clear tissue difference in the intronic ER with testis having higher expression than the other two tissues. Figure 7C,D shows different patterns between the intronic and exonic ERs where in the exonic ER the expression is lowest in the heart, higher in liver and slightly higher at the testis. However in the intronic ER, liver is the tissue that has the lowest expression. These results suggest that expression at unannotated sequence could have biological relevance beyond local annotated exonic sequence.

**Figure 7.**
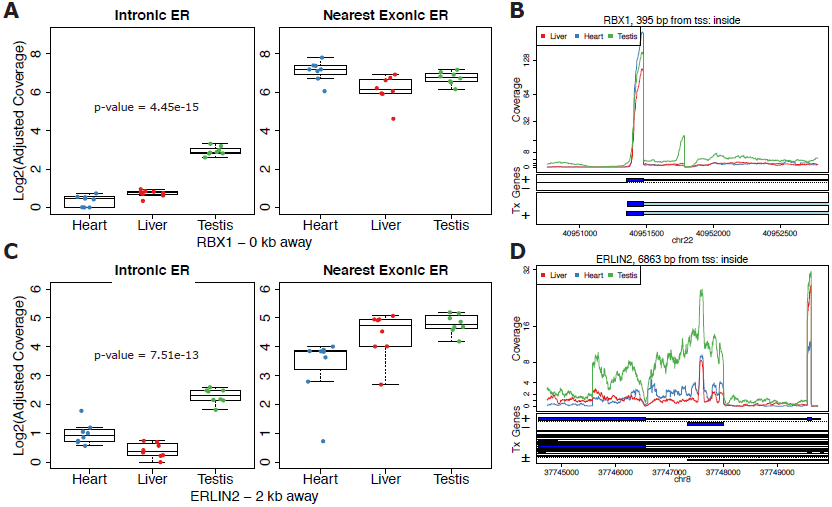
Differential expression on strictly intronic expressed regions adjusting for expression on the nearest strictly exonic ER. Boxplots (**A** and **C**) and region coverage plots (**B** and **D**) for two strictly intronic ERs showing differential expression signal adjusting for the nearest exonic ER. Boxplots show the log_2_ adjusted coverage for the strictly intronic ERs by tissue with the corresponding boxplot for the nearest strictly exonic ERs. The p-value shown is for the differential expression between tissues on the intronic ERs conditional on the expression values for the nearest exonic ERs. The distance to the nearest strictly exonic ER and the gene symbol are shown below. The region coverage plots are centered at the strictly intronic ER with the neighboring 2kb and 5kb for **C** and **D** respectively. **A**,**B** Expression on the exonic ER is fairly similar between the groups but different on the intronic ER. **C**,**D** Expression on the exonic ER has an increasing pattern from heart to liver to testis but has a different pattern on the intronic ER.

### 3.8 Simulation results

We lastly performed a simulation study to evaluate the statistical properties of `derfinder` with and without complete annotation. To compare `derfinder` against feature-level alternatives, we simulated reads for 2 groups, 10 samples in total (5 per group) with ⅙ of the transcripts having higher and ⅙ lower expression in group 2 versus group 1 at fold changes of 2x and ½x respectively. Reads were simulated from chromosome 17 using `polyester` [26] with the total number of reads matching the expected number given paired-end library with 40 million reads (Methods Section 2.4.3). We used `HISAT` [27] to align the simulated reads and summarized them using either featureCounts from the `Rsubread` package [13] or `StringTie` [23] and performed the statistical tests on the resulting coverage matrices using `DESeq2` and `ballgown` [22] respectively. We performed the `ballgown` statistical test at the exon-level as well as the transcript-level. We performed the feature-level analyses using the complete annotation and with an annotation set missing 20% randomly selected transcripts (8.28% unique exons missing). We then used `derfinder` to find the ERs from the same `HISAT` alignments as well as from `Rail-RNA` [24] output and performed the statistical test with `DESeq2`. For all statistical tests we controlled the FDR at 5% and we repeated the simulation three times.

Table 1 shows the range of the empirical power, false positive rate (FPR) and false discovery rate (FDR) for all these methods based on the three simulation replicates. `derfinder`’s expressed region approach resulted in overlapping empirical power ranges to the exon-level methods that are supplied the complete annotation. The exon-level methods had a 18% to 23% loss in power when using the incomplete annotation set compared to the complete set even though only 8.28% of the unique exons were missing. `derfinder`, being annotation-agnostic, does not rely on having the complete annotation but did show increased FPR and FDR compared to the exon-level methods. We recommend performing sensitivity analyses of the cutoff parameter used for defining ERs or the FDR control in the statistical method used to determine which ERs are differentially expressed (i.e. DERs). Transcript-level analyses had the lowest FPR and FDR but also the lowest power. Note that we only performed transcript expression quantification with `StringTie` and did not use the data to determine new transcripts. Doing so resulted in a much larger transcript set than originally present in the data: 3,900 in the original set versus 15,920 (average for the three replicates using the complete annotation). Supplementary Section 1.5.1 shows the results when using `edgeR`-robust for performing the statistical tests. Identifying ERs uses computational resources and runs in similar time to summarization steps required for the exon-level pipelines used in this simulation (Supplementary Section 1.5.2) and is the fastest when using BigWig files such as those produced by `Rail-RNA`. These results suggest that the `derfinder` approach performs well when differentially expressed features overlap known annotation and appear in unannotated regions of the genome.

**Table 1.**
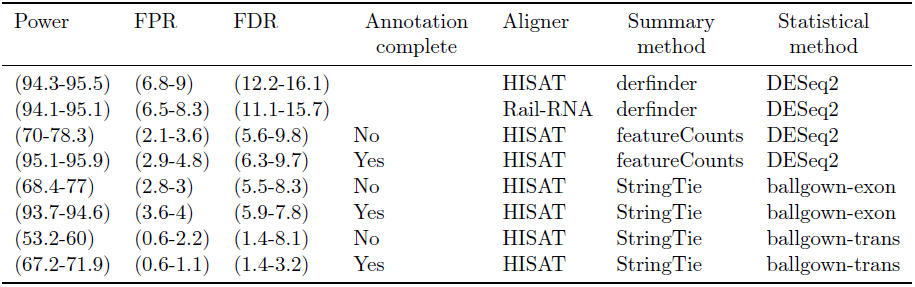
Minimum and maximum empirical power, false positive rate (FPR) and false discovery rate (FDR) observed from the three simulation replicates for each analysis pipeline. `ballgown` analyses were done at either the exon or transcript levels. Pipelines that rely on annotation were run with the full annotation or with 20% of the transcripts missing (8.28% exons missing). Count matrices were analyzed with `DESeq2` and `edgeR`-robust (Supplementary Table S2).

| Power | FPR | FDR | Annotation complete | Aligner | Summary method | Statistical method |
| --- | --- | --- | --- | --- | --- | --- |
| (94.3-95.5) | (6.8-9) | (12.2-16.1) |  | HISAT | derfinder | DESeq2 |
| (94.1-95.1) | (6.5-8.3) | (11.1-15.7) |  | Rail-RNA | derfinder | DESeq2 |
| (70-78.3) | (2.1-3.6) | (5.6-9.8) | No | HISAT | featureCounts | DESeq2 |
| (95.1-95.9) | (2.9-4.8) | (6.3-9.7) | Yes | HISAT | featureCounts | DESeq2 |
| (68.4-77) | (2.8-3) | (5.5-8.3) | No | HISAT | StringTie | ballgown-exon |
| (93.7-94.6) | (3.6-4) | (5.9-7.8) | Yes | HISAT | StringTie | ballgown-exon |
| (53.2-60) | (0.6-2.2) | (1.4-8.1) | No | HISAT | StringTie | ballgown-trans |
| (67.2-71.9) | (0.6-1.1) | (1.4-3.2) | Yes | HISAT | StringTie | ballgown-trans |

## 4 Discussion

Here we introduced the `derfinder` statistical software for performing genome-scale annotation-sagnostic RNA-seq differential expression analysis. This approach utilizes coverage-level information to identify differentially expression regions (DERs) at the expressed region or single base-levels, and then generates useful summary statistics, visualizations and reports to further inspect and validate candidate regions.

The reduced dependence on the transcriptome annotation permits the discovery of novel regulated transcriptional activity, such as the expression of intronic or intergenic sequences, which we highlight in publicly available RNA-seq data and our previous `derfinder` application [20]. As shown with a subset of GTEx, strictly intronic ERs can differentiate tissues when adjusting for the expression from the nearest exonic expressed region, suggesting that some intronic DERs may represent signal beyond run-off transcription. Furthermore, the structure of DERs across a given gene can permit the direct identification of differentially expressed transcripts (e.g. Figure 2C), providing useful information for biologists running validation experiments. Lastly, this software and statistical approach may be useful for RNA-seq studies on less well-studies species, where transcript annotation is especially likely to be incomplete.

The software pipeline, starting with BAM or BigWig files, and ending with lists of DERs, reports, and visualizations, runs at comparable speeds to existing RNA-seq analysis software. Given the appropriate computing resources, `derfinder` can scale to analyze studies with several hundred samples. For such large studies, it will be important to correct for batch effects and potentially expand `derfinder`’s statistical model for base-level covariates. This approach provides a powerful intermediate analysis approach that combines the benefits of feature counting and transcript assembly to identify differential expression without relying on existing gene annotation.

## 7 Author’s contributions

AEJ, JTL, RAI conceived the software. LCT wrote the software under the supervision of JTL and AEJ. LCT analyzed the data with the supervision of JTL and AEJ. AN, CW and BL helped with the GTEx data analysis. All authors contributed to writing the paper.

## 8 Acknowledgments

The Genotype-Tissue Expression (GTEx) Project was supported by the Common Fund of the Office of the Director of the National Institutes of Health. Additional funds were provided by the NCI, NHGRI, NHLBI, NIDA, NIMH, and NINDS. Donors were enrolled at Biospecimen Source Sites funded by NCI/SAIC-Frederick, Inc. (SAIC-F) subcontracts to the National Disease Research Interchange (10XS170), Roswell Park Cancer Institute (10XS171), and Science Care, Inc. (X10S172). The Laboratory, Data Analysis, and Coordinating Center (LDACC) was funded through a contract (HHSN268201000029C) to The Broad Institute, Inc. Biorepository operations were funded through an SAIC-F subcontract to Van Andel Institute (10ST1035). Additional data repository and project management were provided by SAIC-F (HHSN261200800001E). The raw data (sequencing reads and phenotype data) used for the analyses described in this manuscript were obtained from SRA accession number phs000424.v6.p1 on 10/07/2015.

## 9 Additional Files

The `derfinder` vignettes detail how to use the software and its infrastructure. The latest versions are available at www.bioconductor.org/packages/derfinder.

The Supplementary Methods and Results describe in more detail the R implementation, the single base-level approach, and the analysis of the *BrainSpan* data set with the single base-level approach. Supplementary file 1 contains the identified candidate single base-level DERs in CSV format (gzip compressed) for the *BrainSpan* data set.

The code and log files detailing the versions of the software used for all the analyses described in this paper is available at the Supplementary Website: leekgroup.github.io/derSupplement.

## Supplementary Methods and Results

This document describes R implementation details of `derfinder`, the single base-level approach, and results from applying the single base-level approach to the *BrainSpan* data set. It also includes the simulation results when performing the statistical tests using `edgeR`-robust [1] instead of *DESeq2* [2].

## 1 Supplementary Results

### 1.1 R implementation

The `derfinder` package can be used for different types of analyses such as DER finding (single base-level and ER-level approaches) as well as creating a feature counts matrix. The overall relationship between these functions is shown in section *Flow charts* subsection *DER analysis flow chart* of the *derfinder users guide* vignette available at www.bioconductor.org/packages/derfinder.

For the single base-level approach, the main function is `analyzeChr()` which makes it easier for users to run this type of analysis. This function is a wrapper for other functions available in `derfinder`, as can be seen section *Flow charts* subsection *analyzeChr() flow chart* of the *derfinder users guide* vignette. It splits the data, calculates the F-statistics, identifies the null regions, and annotates them.

The expressed regions (ERs) approach is described in section *Flow charts* subsection *regionMatrix() flow chart* of the *derfinder users guide* vignette. This type of analysis requires fewer functions, as the user only needs to load the data and then identify the ERs with the `regionMatrix()` function. The *regionMatrix() flow chart* shows which other functions are internally used by `regionMatrix()` that filter the coverage by using a mean cutoff, identify the regions, and produce the region-level count matrix. The function `railMatrix()` is optimized for identifying ERs from BigWig files, specially those created with `Rail-RNA` (DOI: 10.1101/019067).

### 1.2 Single base-level statistical test

A single base-level resolution analysis in `derfinder` starts with read alignment and coverage calculation as done in the ER-level approach. Next, a standard differential expression analysis is performed at each base by comparing nested null and alternative linear models using an F-statistic. The statistical models may include adjustments for confounders such as library size [3], demographic variables, and batch effects [4].

Once an F-statistic is calculated at each base, we identify differentially expressed regions (DERs) using a “bump hunting” approach [5]. First we find candidate DERs by identifying regions of the genome where the base-level F-statistics pass a genome-wide threshold (Figure S1 with *BrainSpan* data set, see Supplementary Section 2.1). We then calculate a summary statistic for each candidate region based on the length of the region and the size of the statistics within the region. To evaluate the statistical significance of these candidate regions, we permute the sample labels and recompute candidate regions and summary statistics. The result is a region-level p-value, which can be adjusted to control the family-wise error rate. Alternatively, the region-level p-values can be adjusted for multiple testing using standard false discovery rate techniques [6, 7].

**Figure S1.**
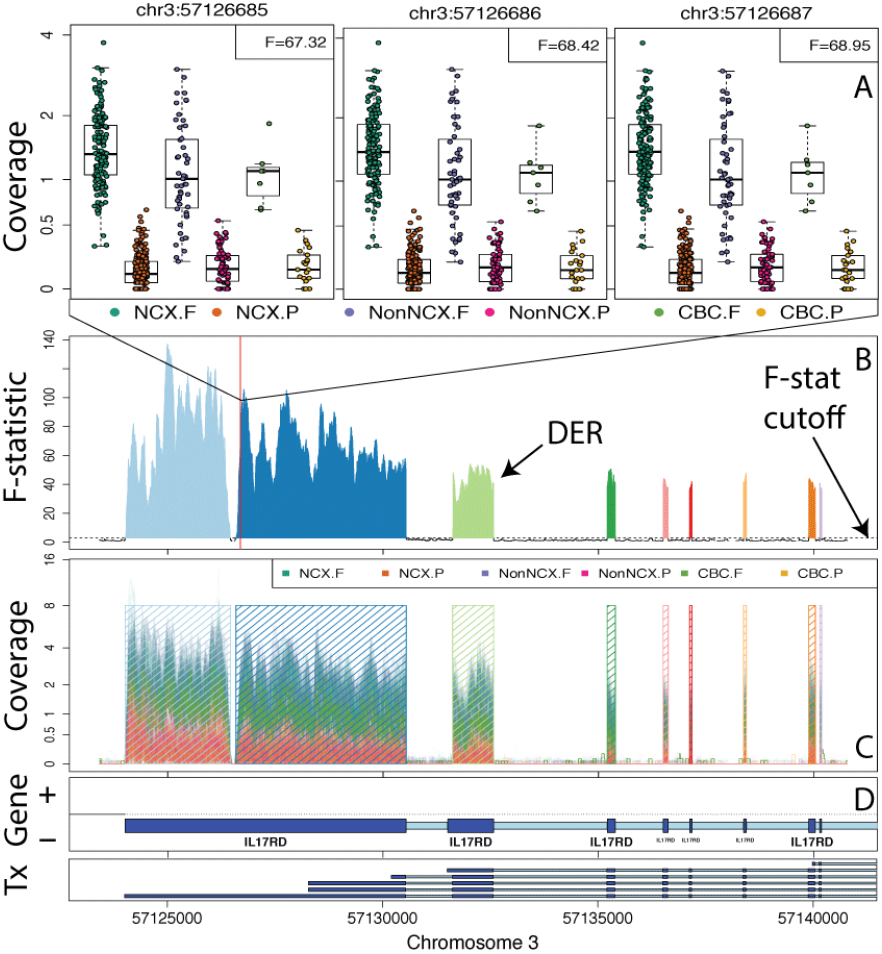
Finding DERs on chromosome 3 with *BrainSpan* data set using six groups. Neocortical regions (NCX: DFC, VFC, MFC, OFC, M1C, S1C, IPC, A1C, STC, ITC, V1C), Non-neocortical regions (NonNCX: HIP, AMY, STR, MD), and cerebellum (CBC) split by whether the sample is from a fetal (F) or postnatal (P) subject. **A** Boxplots for three specific bases. **B** F-statistics curve with regions passing the F-stat cutoff marked as candidate DERs. **C** Raw coverage curves superimposed with the candidate DERs. **D** Known exons (dark blue) and introns (light blue) by strand. The third DER matches the shorter version of the second exon shown in the *Tx* track.

### 1.3 Differential expression in the developing human brain via expressed region-level analysis

Figure S2 complements Figure 5 with the results of performing principal component analysis of ERs found in the *BrainSpan* data set given the known annotated elements they overlap with. The results are consistent regardless of the type of ERs under study.

**Figure S2.**
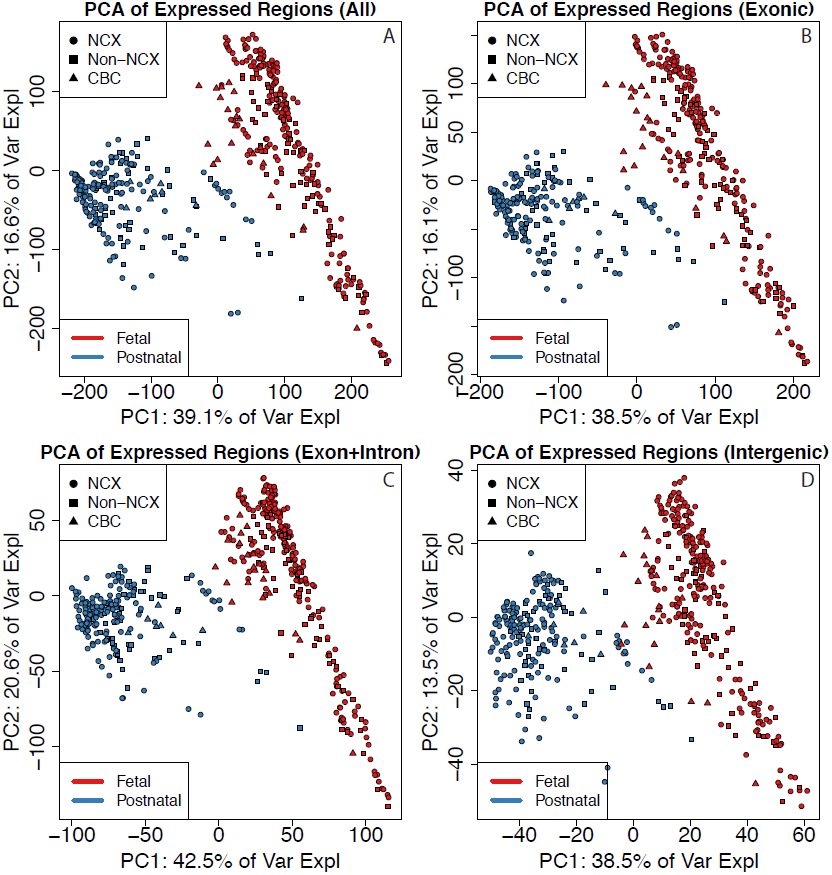
Principal components analysis reveals clusters of samples in the BrainSpan data set. First two principal components (PCs) with samples colored by sample type (F: Fetal or P: Postnatal) and shape given by brain region using all ERs (top left), strictly exonic ERs (top right), ERs overlapping exons and introns (bottom left) and strictly intergenic ERs (bottom right).

### 1.4 Differential expression in the developing human brain via single base-level analysis

At the single base-level, we identified 113,691 genome-wide significant DERs (FWER < 5%) with the same statistical models used with the ER-level analysis described in the main text. These resulting single base-level DERs largely distinguished the fetal and postnatal samples representing the first principal component and 49.4% of the variance of the mean coverage levels within the DERs (Figure S3). The most significant DERs map to genes previously implicated in development, and contained many of the DERs we previously identified in the frontal cortex in 36 independent subjects [8]. For example, 59% of our previously published 50,650 developmental DERs (and 72.6% in the 10,000 most significant) in the frontal cortex overlapped these DERs identified in the *BrainSpan* data set. The potential lack of overlap may be explained by unmodeled artifacts as there appear to be clusters in the principal components calculated on the base resolution data (Figure S3, left panel).

**Figure S3.**
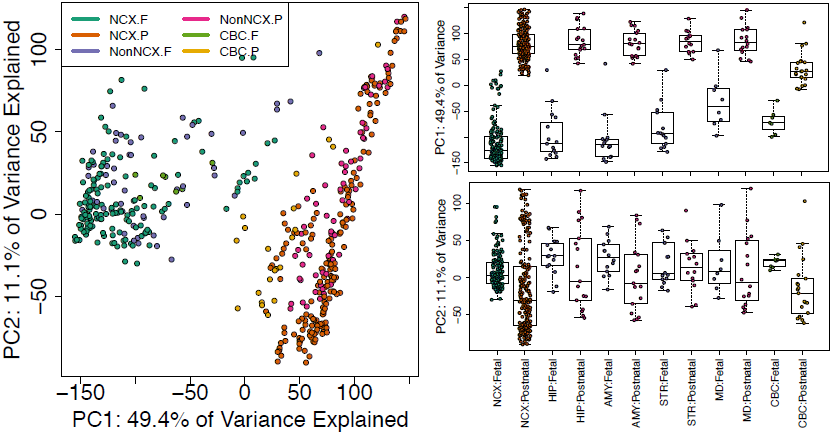
Principal components analysis reveals clusters of samples in the BrainSpan data set. (Left) First two principal components (PCs) with samples colored by sample type (F: Fetal or P: Postnatal) and shape given by brain region. (Right) Boxplots for PCs 1 and 2 by brain region (NCX: neocortex, HIP: hippocampus, AMY: amygdala, STR: striatum, MD: thalamus, CBC: cerebellum) and sample type with non-neocortex brain decomposed into its specific regions.

While the majority (68.1%) of single base-level DERs overlap exclusively exonic sequence using Ensembl database v75, we find that a fraction (22.2%) of the single base-level DERs map to sequence previously annotated as non-exonic (e.g. solely intronic or intergenic). The proportion of exonic sequence is higher than our previous analyses in the frontal cortex [8]. When the single base-level DERs are stratified by brain region and developmental period with the highest expression levels (Table S1), we find the highest degree of unannotated regulation in the cerebellum, the brain region with the largest degree of region-specific genes in a previous analyses [9]. The majority of DERs, regardless of their annotation, are most highly expressed in fetal life, particularly within the neocortex, hippocampus, and amygdala. Non-exonic expression might be due to incomplete transcript annotation in reference databases, background expression, or previously undetected artifacts.

**Table S1.**
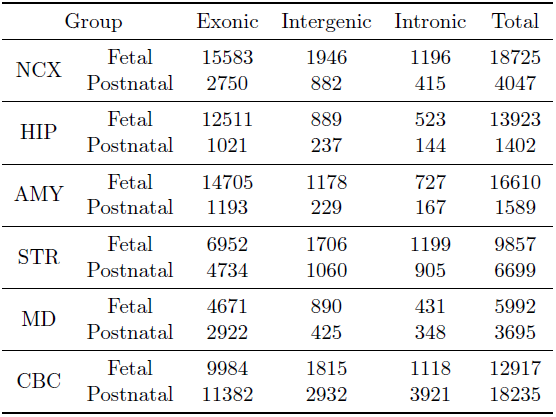
Classification of single base-level DERs in the *BrainSpan* project. For each statistically significant DER, we identified the developmental period and region with the highest average expression levels, stratified by annotation relative to the Ensembl gene database. NCX: neocortex, HIP: hippocampus, AMY: amygdala, STR: striatum, MD: thalamus, CBC: cerebellum. Region assignment is prioritized by exon > intron > intergenic.

### 1.5 Simulation analysis

#### 1.5.1 Simulation results with edgeR-robust

Table S2 shows the empirical power, false positive rate (FPR) and false discovery rate (FDR) for the different analysis pipelines that result in a count matrix which we analyzed with `edgeR`-robust [10] while controlling the FDR to 5%. The observed power is slightly higher than the corresponding results using `DESeq2` [2]. The observed FPR and FDR with `edgeR`-robust are higher than in the `DESeq2` results, with overlapping ranges for the `derfinder` analyses and non-overlapping ones when summarizing the data with `featureCounts` [1].

**Table S2.**
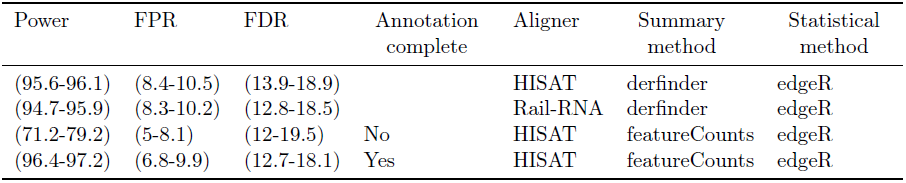
Simulation results for pipelines that used edgeR-robust for the statistical tests. Minimum and maximum empirical power, false positive rate (FPR) and false discovery rate (FDR) observed from the three simulation replicates for each analysis pipeline that resulted in a count matrix analyzed with `edgeR`-robust. `ballgown` analyses were done at either the exon or transcript levels. Pipelines that rely on annotation were run with the full annotation or with 20% of the transcripts missing (8.28% exons missing).

#### 1.5.2 Timing and computational resources used

Table S3 shows a summary of the computational resources used for the different pipelines used in the simulation as well as the time for running them. In general, the maximum memory per core is low (most are below 3.2 GB) regardless of the analysis step. The exception is alignment with `Rail-RNA` because of how our computing cluster measures memory usage: it artificially increases when processes spawn shared-many sub-processes by counting more than once the memory used by shared objects. Time-wise all analysis steps except for alignment take only 11 minutes at most. Notably, the ER-level approach is much faster with `Rail-RNA` output than with `HISAT` output. This is because `derfinder` can load the data much faster from BigWig files than from BAM alignment files and the `railMatrix` has been optimized for the BigWig files that `Rail-RNA` produces. In this particular simulation, `Rail-RNA` is slower than `HISAT` for aligning reads, but this is expected since `Rail-RNA` is better suited at analyzing larger data sets in the cloud and decreasing false positives when determining new splice junctions. This is reflected on Table 1 and S2 with slightly reduced FPR and FDR when using `Rail-RNA` compared to `HISAT`. The timing results for each computing job are available in the Supplementary Website.

**Table S3.** Summary of computing resources required for each analysis step for the different simulation pipelines. This table shows the maximum memory (GB) per core, the time in minutes to run the analysis with all jobs running sequentially and the maximum number of cores used in any step of the simulation analysis for the different pipelines. Note that the ERs (HISAT), the feature-level counts and ballgown pipelines rely on HISAT alignments.

| Max memory<br>by core (GB) | Time (minutes) | Peak cores | Pipeline | Analysis step |
| --- | --- | --- | --- | --- |
| (2.8-3.1) | (2.1-3.2) | 10 | ER-level (**Rail-RNA**) | Align prep |
| (32.8-39.1) | (137.6-218.4) | 10 | ER-level (**Rail-RNA**) | Align |
| (3.2-3.2) | (47.2-72.1) | 40 | **HISAT** | Align |
| (1.4-1.4) | (1.5-1.5) | 1 | ER-level (**Rail-RNA**) | Summarize |
| (0.6-0.6) | (1.3-1.9) | 1 | ER-level (**Rail-RNA**) | Statistical tests |
| (0.8-0.8) | (5-7) | 4 | ER-level (**HISAT**) | Summarize |
| (0.6-0.6) | (1.5-1.9) | 1 | ER-level (**HISAT**) | Statistical tests |
| (2.2-2.2) | (1.6-1.6) | 8 | Feature counts | Summarize |
| (0.6-0.6) | (3.7-5.3) | 2 | Feature counts | Statistical tests |
| (2.1-2.1) | (8.7-11) | 80 | StringTie-Ballgown | Summarize |
| (0.7-0.7) | (0.7-0.8) | 2 | StringTie-Ballgown | Statistical tests |

## 2 Supplementary Methods

### 2.1 single base-level derfinder

The single base-level approach implemented in `derfinder` requires two models. The alternative model (1) contains an intercept, the primary covariate of interest, and optionally adjustment variables. The primary variable can be as simple as a case-control variable or a more complicated model including smoothing functions (e.g. splines) over time. The adjustment variables can include a library size normalization factor for raw data and optionally other potential confounders like age, sex, and batch variables. There are different library size normalization factors you can consider using and `derfinder` implements a version in the `sampleDepth` function based on Paulson et. al [11].

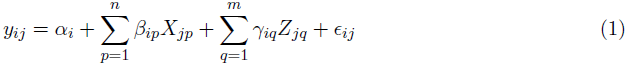

In both models *y_ij_* is the scaled log_2_ base-level coverage for genomic position *i* and sample *j*. That is, *y_ij_* = log_2_ (coverage_*ij*_ + scaling factor). The model is completed by the *n* group effects *β*_i_, *m* adjustment variable effects *γ*_i_ and potentially correlated measurement error *ϵ*. The null model (2) is nested within model (1) and contains only the intercept and adjustment variables.

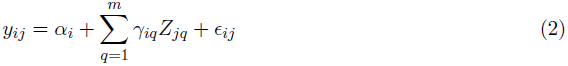

`derfinder` uses a fixed design matrix, testing the same hypothesis at every base. This permits fast vectorized differential expression analysis. At each base we compute a moderated F-statistic

[12] of the form in equation (3), where RSS0_*i*_ and RSS1_*i*_ are the residual sum of squares of the null and alternative models for base *i*. Furthermore, df_0_ and df_1_ are the degrees of freedom for the null (2) and alternative (1) models respectively, *n* is the number of samples, and an offset can be used for smaller experiments to shrink large F-statistics that may be driven by few biological replicates that cluster tightly.

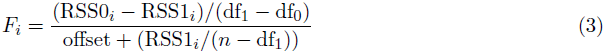

We then perform “bump hunting” adapted to `Rle` objects in order to identify candidate DERs, *R_k_*. Candidate DERs are defined as contiguous sets of bases where *F_i_* > *T* for a fixed threshold *T*. We then calculate an “area” statistic for each candidate DER which is the sum of the F-statistics above the threshold within the region: *S_k_* = Σ_*j*∈*R_k_*_ *F_j_* (Figure S1B). We have previously applied this approach to identify local differentially and variably methylated regions and more long range changes in methylation [5, 13, 14]. One key difference compared to previous implementations in DNA methylation data is that we do not explicitly smooth the F-statistics, allowing for precise discovery of intron-exon boundaries in the data (Figure S1C).

Permutation analysis generates statistical significance for each of these candidate DERs by permuting the sample labels, re-calculating the F-statistics, identifying null candidate regions and region-level statistics in this permuted data set, and then calculating empirical p-values and/or directly estimating the family-wise error rate (FWER) [5]. Alternatively, the empirical p-values can be adjusted to control the false discovery rate (FDR) via qvalue [6].

### 2.2 Data Processing: BrainSpan data

For the single base-level analysis, we used a scaling factor of 1 and chose the F-statistic cutoff *T* such that *P*(*F* > *T*) = 10^−6^. We used the same alternative model described for the expressed region analysis in the main text. We compared the alternative model to an intercept-only model, and identified DERs using the single base-level analysis. We then calculated the mean coverage for each significant single base-level DERs in each sample, resulting in a mean coverage matrix (DERs by samples), and we performed principal component analysis (PCA) on this log_2_-transformed matrix (after adding an offset of 1), which were subsequently plotted in Figure S3.

